# Bone Lymphatic Endothelial Cells Form a Distinct Skeletal Population and Drive Skeletal Repair

**DOI:** 10.64898/2026.08.06.743157

**Authors:** Sanyam Jain, Jie Li, Jiayuan Zhang, He Cai, Yufei Wu, Yang Yang, Minghan Ye, Makarand V Risbud, Junyu Chen, Anjali P. Kusumbe

## Abstract

Bone regeneration relies on specialized vascular niches, yet the contribution of lymphatic vessels across distinct skeletal sites remains poorly understood. Here, we identify bone lymphatics as an essential component of the regenerative microenvironment in the mandible and long bones. We demonstrate that bone lymphatic endothelial cells (LECs) constitute a specialized endothelial population that is transcriptionally and spatially distinct from periosteal LECs. During skeletal repair, bone LECs reactivate a regenerative transcriptional programme, and promote mandibular and fracture healing. In osteonecrosis of the jaw and periodontitis, bone lymphatic-associated signalling is disrupted, identifying impaired lymphatic function as a shared feature of mandibular disease. Therapeutic activation of VEGFC–FLT4 signalling during injury or mandibular disease restores lymphangiogenesis, enhances osteogenesis, and markedly improves bone regeneration. Together, our findings advance the paradigm-shifting role of bone lymphatics positive regulators of bone regeneration and identify lymphatic activation as a promising therapeutic strategy to enhance bone regeneration in mandibular diseases.

## Introduction

Bone regeneration is orchestrated by specialized microenvironments that coordinate vascular, immune and skeletal cell interactions to maintain tissue homeostasis and repair^1-7^. The skeletal vasculature plays a central role in this process by providing angiocrine signals that regulate osteogenesis, hematopoiesis and stem cell function^8-14^. While blood vessels have been extensively studied as regulators of bone biology, the contribution of lymphatic vessels to skeletal homeostasis has only recently begun to emerge^15-18^. Understanding how different vascular compartments cooperate to regulate bone regeneration is therefore essential for developing regenerative therapies for skeletal disease.

Lymphatic vessels are increasingly recognized as active regulators of tissue repair rather than passive conduits for fluid drainage. Beyond maintaining tissue fluid balance and immune cell trafficking^19-21^, lymphatic endothelial cells (LECs) produce instructive signals that regulate inflammation, stem cell activity and tissue regeneration in multiple organs^15,22-25^. In the skeleton, accumulating evidence has fundamentally changed the long-standing view that mineralized bone lacks lymphatic vasculature^15,17,26-28^. Studies from several independent groups have identified the role of lymphatic vessels during the bone repair and regeneration^16,17,28-36^. These studies demonstrate that lymphatic dysfunction contributes to age-associated bone loss^17^, inflammatory osteolysis^37,38^ and impaired fracture repair^28,39,40^, whereas enhancing lymphangiogenesis promotes bone formation, limits osteoclast activation and supports regenerative immune responses^17,18,25,41-45^. Together, these findings position bone lymphatics as active regulators of skeletal homeostasis rather than passive anatomical structures.

The mandible represents a unique skeletal tissue in which efficient vascular coordination is particularly critical. Unlike long bones, mandibular bone is continuously exposed to mechanical loading, microbial challenge and repetitive injury associated with mastication and dental procedures^46-49^. Consequently, disorders such as periodontitis and osteonecrosis of the jaw (ONJ) are characterized by chronic inflammation, defective vascular remodeling and failure of bone regeneration^49-53^. Although impaired blood vessel function has been implicated in these disorders^54,55^, the contribution of lymphatic vessels to mandibular homeostasis and repair remains poorly understood. Whether disruption of lymphatic signaling represents a common mechanism underlying these clinically distinct diseases has not been determined.

Several key questions therefore remain unresolved. It is unknown whether bone lymphatics actively regulate mandibular regeneration through tissue-specific signaling mechanisms, whether their dysfunction is a conserved feature of mandibular disease, and whether therapeutic activation of lymphatic pathways can restore bone repair. Addressing these questions is particularly important because lymphatic endothelial cells are increasingly recognized as sources of regenerative lymphangiocrine factors capable of coordinating vascular remodeling, immune responses and tissue regeneration.

Here, we define bone lymphatics as an essential component of the mandibular regenerative microenvironment. By integrating spatial molecular profiling, single-cell transcriptomics, three-dimensional tissue analyses and functional disease models, we demonstrate that mandibular lymphatic endothelial cells adopt a distinct regenerative transcriptional program during bone repair. We further show that lymphatic-associated signaling is consistently disrupted across experimental and human mandibular diseases, including periodontitis and osteonecrosis of the jaw. Finally, we demonstrate that therapeutic activation of VEGFC–FLT4 signaling restores lymphatic function, enhances vascular and osteogenic regeneration, and markedly improves healing in experimental osteonecrosis of the jaw. Together, our findings identify bone lymphatics as central regulators of mandibular regeneration and establish the lymphatic vasculature as a promising therapeutic target for craniofacial bone repair.

## Results

### Transcriptomic analyses distinguish bone lymphatic endothelial cells from periosteal lymphatics and identify bone LECs as a distinct skeletal cell population

Analysis of publicly available periosteal single-cell RNA-sequencing datasets demonstrates that periosteal LECs are unlikely to represent the source of the LEC population identified in bone single-cell RNA-sequencing datasets. Not all periosteal datasets contain a discrete LEC cluster. Periosteal stromal datasets prepared from pooled tissues from several mice frequently showed no identifiable LEC cluster (e.g., GSE272612 and GSE249528) and contained only rare or undetectable Prox1-expressing cells (Fig. 1a). In one dataset (GSM8123674), a small number of LECs (15 cells from 3 mice) could be recovered; however, these cells remained scarce, and across datasets, the *Prox1*-positive cells often lacked robust *Lyve1* co-expression or failed to resolve into a well-defined canonical lymphatic endothelial cluster (Fig. 1a). Collectively, these observations indicate that the periosteum harbors too few LECs to account for the reproducible *Prox1*-positive LEC population consistently identified across independent bone and bone marrow datasets.

**Figure 1.**
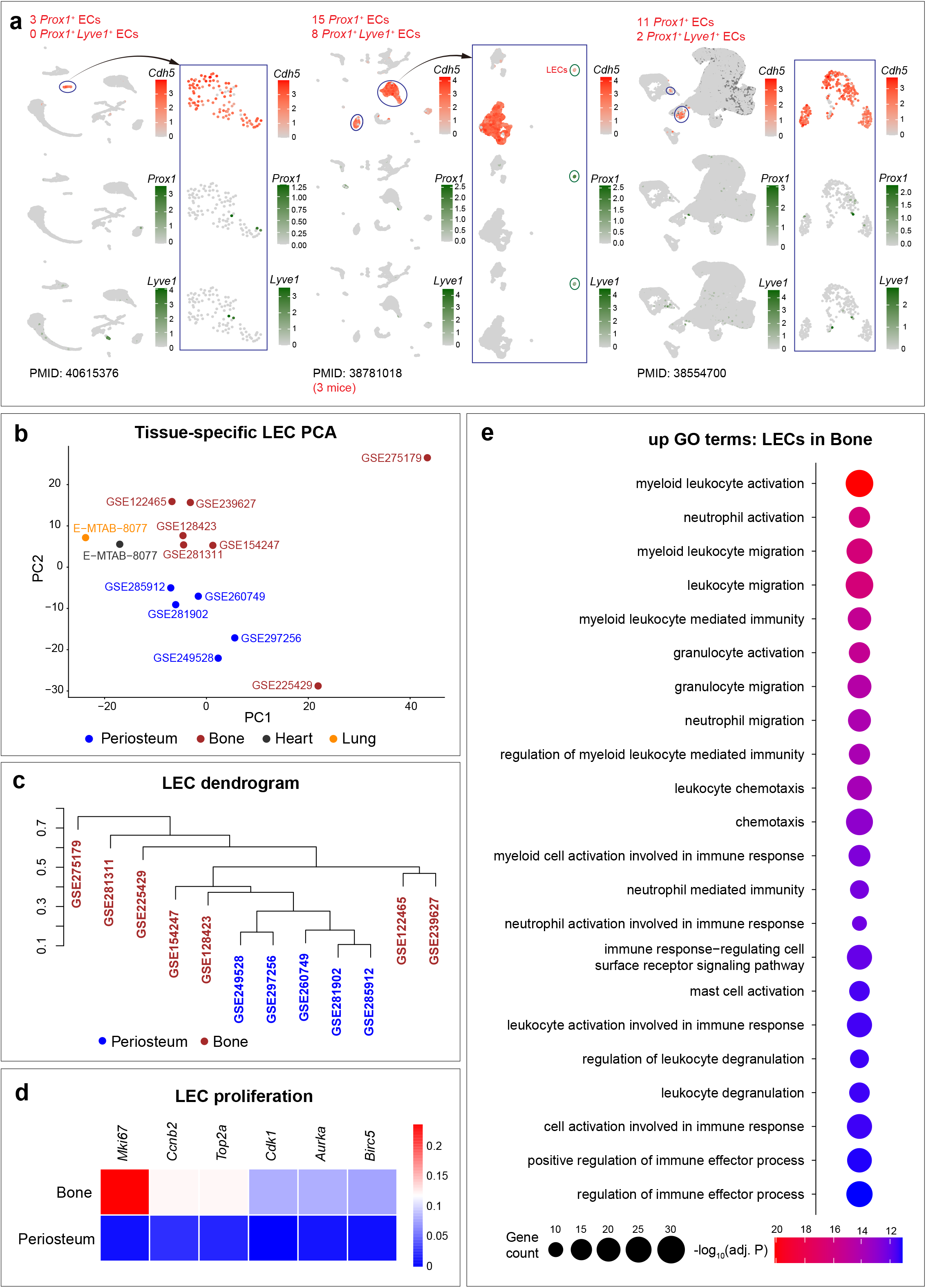
Distinct molecular profile of bone and periosteal lymphatic endothelial cells (LECs). **a** Feature plots showing expression patterns of *Cdh5*, *Prox1* and *Lyve1* in GSE272612 (GSM8406759-61) from PMID: 40615376, GSE260749 (GSM8123764) from PMID: 38781018 and GSE249528 (GSM7949347-50) from PMID: 38554700. **b** Principal component analysis (PCA) of transcriptomic profiles of LECs from periosteum, bone, heart and lung tissues. **c** Dendrogram obtained from hierarchal clustering of LECs from bone and periosteum datasets. **d** Heatmap showing differential expression of proliferation-related genes in bone vs periosteal LECs. **e** Gene ontology (GO) term enrichment on genes upregulated in bone LECs compared to periosteal LECs.

We next investigated whether periosteal lymphatic cells share a common transcriptional identity with bone LECs. To address this, we integrated periosteal and bone datasets and performed principal component analysis together with differential gene expression analysis between LECs of the two tissues. Rather than overlapping with bone LECs, periosteal lymphatic cells segregated into a distinct transcriptional space (Fig. 1b), demonstrating that the two populations represent molecularly distinct endothelial identities. This is also evidenced by the closeness of periosteal LECs from multiple independent datasets in hierarchical clustering (Fig. 1c). Importantly, principal component analysis showed that periosteal lymphatic endothelial cells clustered more closely with lymphatic endothelial cells from classical soft tissues, including lung and heart, whereas bone-derived LECs occupied a separate transcriptional space (Fig. 1b). These findings indicate that bone LECs possess a transcriptional identity that is distinct from both periosteal and conventional soft-tissue lymphatic endothelium. Differential gene expression analysis further supported this distinction by revealing markedly different transcriptional program between bone and periosteal LECs (Fig. 1d). Bone LECs preferentially expressed genes associated with *Mki67*, *Ccnb2*, and *Top2a*, whereas periosteal LECs exhibited a molecular signature closely resembling that of peripheral soft-tissue lymphatics (Fig. 1d). The concordant separation observed across both unsupervised dimensionality reduction and differential expression analyses demonstrates that periosteal lymphatic endothelial cells are transcriptionally distinct from bone LECs and therefore are unlikely to represent their cellular source. Gene ontology analysis of genes upregulated in bone LECs highlighted enrichment for myeloid leukocyte activation, neutrophil and granulocyte activation/migration, leukocyte chemotaxis, and related immune effector processes (Fig. 1e). The concordant separation observed across unsupervised dimensionality reduction, hierarchical clustering, differential expression, and functional enrichment demonstrates that periosteal LECs are both quantitatively insufficient and transcriptionally distinct from bone LECs. Thus, periosteal LECs cannot account for the bone-associated LEC population. Instead, bone-associated LECs constitute a distinct skeletal lymphatic endothelial population with a unique transcriptional identity, indicating that they represent a bona fide component of the bone microenvironment rather than cells originating from the periosteum.

### Bone lymphatic endothelial cells expand and acquire a proliferative, metabolically active state following fracture

Spatial transcriptomic analysis of fractured bone identified *Prox1*^+^ endothelial cells embedded within the bone matrix, suggesting that bone lymphatic endothelial cells (LECs) respond dynamically to skeletal injury^25^. To further define the behavior of bone LECs during regeneration, we analyzed publicly available single-cell RNA-sequencing datasets comprising 24 enriched stromal cell samples from fractured and uninjured bones, integrated using Harmony (Fig. 2a). Bone LECs, identified by expression of *Prox1*, *Lyve1*, and *Flt4*, expanded markedly following fracture, increasing from 1.13% of endothelial population in uninjured bone to 3.16% after injury, whereas arterial and sinusoidal endothelial cell populations were proportionally reduced (Fig. 2a). Consistent with their expansion, fracture-associated LECs exhibited a pronounced proliferative signature, characterized by increased expression of cell-cycle regulators including *Top2a*, *Mki67*, *Cdc20*, *Birc5*, and *Ccnb2*, demonstrating active lymphangiogenesis during skeletal repair (Fig. 2b).

**Figure 2.**
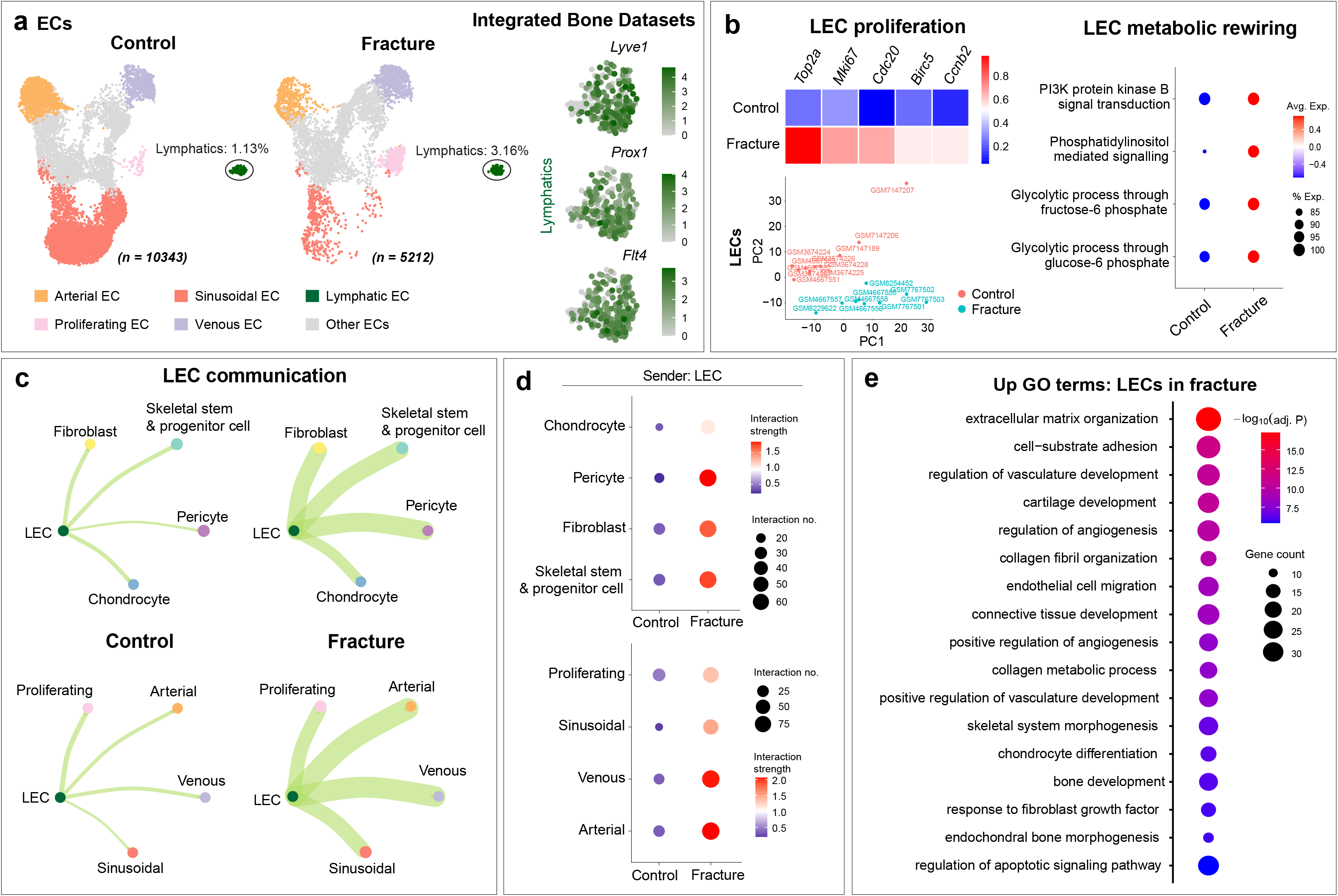
Bone lymphatic endothelial cells expand and acquire pro-regenerative transcriptional program following fracture. **a** UMAPs showing different bone endothelial sub-populations under control vs fracture conditions. Feature plots showing expression of *Lyve1*, *Prox1* and *Flt4* in lymphatic endothelial cells (LECs). **b** Heatmap showing differential expression of proliferation-related genes in bone LECs under control vs fracture conditions. Principal component analysis (PCA) of transcriptomic profiles of bone LECs under control vs fracture conditions. Dot plot showing gene-ontology (GO) term scores bone LECs under control vs fracture conditions. **c** Network plots showing predicted communication of bone LECs with skeletal stem and progenitor cells, pericytes and blood-vascular endothelial cell sub-types in control vs fracture conditions. **d** Dot plots indicating number and strength of interactions from bone LECs to mesenchymal and blood-vascular endothelial cell sub-types under control vs fracture conditions. **e** GO term enrichment on genes upregulated in bone LECs under fracture vs control conditions.

Principal component analysis further showed that fracture-associated proliferating LECs segregated into a distinct transcriptional state from homeostatic LECs (Fig. 2b), indicating extensive injury-induced cellular reprogramming. Gene Ontology term scoring further revealed significant enrichment in glycolytic metabolism, consistent with the metabolic shift accompanying endothelial activation and proliferation during tissue regeneration (Fig. 2b).

### Fracture reprograms bone lymphatic endothelial cells into regenerative signaling hubs

To determine whether activated LECs also acquire specialized regenerative functions, we next performed CellChat analysis. Fracture substantially increased both the number and strength of signaling interactions between LECs and multiple skeletal cell populations, including skeletal stem and progenitor cells (SSPCs), fibroblasts, pericytes, chondrocytes, and vascular endothelial cells (Fig. 2c,d). Notably, the strongest increase in communication occurred between LECs and SSPCs, suggesting that lymphatic endothelial cells directly engage skeletal progenitors during bone regeneration.

To identify molecular pathways underlying these interactions, Gene ontology analysis of upregulated genes in fracture-associated LECs demonstrated significant enrichment of biological processes involved in bone development, skeletal system morphogenesis, endochondral ossification, extracellular matrix organization, collagen fibril organization, collagen metabolism, endothelial cell migration, regulation of angiogenesis, connective tissue development, and chondrocyte differentiation, indicating that activated LECs acquire a regenerative transcriptional program closely linked to bone formation and tissue remodeling (Fig. 2e).

### Activated bone lymphatic endothelial cells acquire a pro-regenerative lymphangiocrine secretome

We next investigated whether these transcriptional changes translated into alterations in the lymphangiocrine secretome. Analysis of secreted factors expressed by fracture-associated LECs revealed robust upregulation of genes encoding extracellular proteins associated with bone development, skeletal morphogenesis, endochondral ossification, extracellular matrix organization, collagen fibril organization, cell-substrate adhesion, and cartilage development (Fig. 3a). Prominent secreted factors included multiple collagens (*Col1a1*, *Col1a2*, *Col3a1*, *Col6a1*), matrix remodeling enzymes (*Mmp2* and *Mmp13*), matricellular proteins (*Thbs1* and *Spp1*/osteopontin), extracellular matrix glycoproteins (*Fn1*), and osteogenic regulators including *Sparc*, supporting the concept that proliferating bone LECs actively promote skeletal repair through regenerative lymphangiocrine signaling (Fig. 3a).

**Figure 3.**
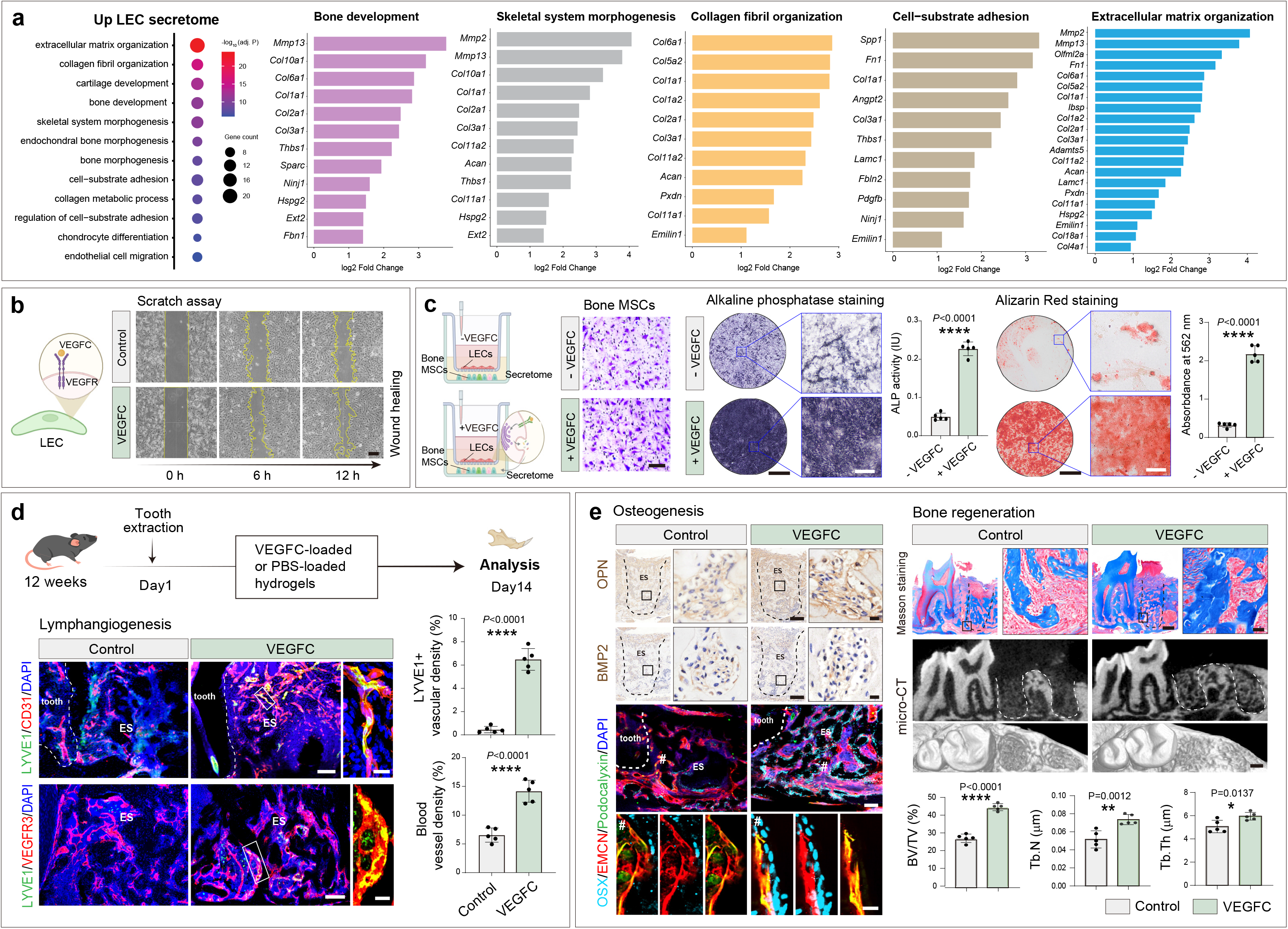
Bone lymphatics orchestrate regenerative signaling to drive bone repair. **a** Gene-ontology (GO) term enrichment on secretome genes upregulated in bone LECs under fracture vs control conditions. Bar plots showing increase in expression of secretome genes belonging to enriched GO terms. **b** Representative images of scratch assay for LECs in presence and absence of VEGFC at 0, 6, and 12 hours. Scale bar: 100 μm. **c** Schematic representing transwell co-culture system with LECs and bone MSCs. Representative images and quantification of alkaline phosphatase staining and Alizarin Red staining of bone MSCs in -VEGFC and +VEGFC groups. *P* value is derived from two-tailed unpaired t tests. **** *P* < 0.0001; *n=5*. Scale bars: 1 mm (black), 50 μm (white). **d** Schematic overview of VEGFC-loaded or PBS-loaded hydrogel treatment after tooth extraction and analysis at day 14. 3D images showing immunolabeling for CD31 and LYVE1. Nuclei: DAPI. Bar graph showing quantification of LYVE1^+^ lymphatic vascular density. *P* value is derived from two-tailed unpaired *t* tests. **** *P* < 0.0001; *n=5*. Scale bars: 100 μm (overview) and 20 μm (insets). **e** Representative images showing immunohistochemical staining for OPN and BMP2, and immunofluorescence staining for OSX, EMCN, and Podocalyxin. Nuclei:DAPI. Representative Masson’s trichrome staining, micro-CT reconstruction images, and quantitative analysis of the micro-CT parameters. *P* value is derived from two-tailed unpaired *t* tests. * *P* < 0.05; ** *P* < 0.01; and **** *P* < 0.0001; *n=5*. ES, extraction socket. Scale bars: for immunostaining images, 150 μm (overview) and 20 μm (insets); for Masson staining 400 μm (overview) and 20 μm (insets).

### VEGFC-activated bone lymphatic endothelial cells promote osteogenic differentiation through lymphangiocrine signaling

To determine whether this injury-induced regenerative program functionally promotes osteogenesis, we next enhanced LEC activation *in vitro*. Because VEGFC is a potent driver of lymphatic endothelial cell proliferation and migration^56^, cultured bone LECs were stimulated with VEGFC. VEGFC markedly accelerated wound closure in scratch assays, confirming enhanced LEC migration and activation (Fig. 3b). LECs were then cultured in the upper chamber of a Transwell co-culture system in the presence or absence of VEGFC, while bone mesenchymal stem cells (MSCs) undergoing osteogenic differentiation were cultured in the lower chamber, allowing communication exclusively through LEC-derived secreted factors without direct cell-cell contact (Fig. 3c). MSCs exposed to VEGFC-activated LECs exhibited significantly increased alkaline phosphatase (ALP) activity, an established marker of early osteoblast differentiation, together with markedly increased Alizarin Red staining, indicating enhanced extracellular matrix mineralization and late-stage osteoblast maturation (Fig. 3c). These findings demonstrate that activated LECs acquire a pro-regenerative lymphangiocrine secretory phenotype that directly stimulates osteogenic differentiation of skeletal progenitor cells.

### Enhancing bone lymphatic activation accelerates skeletal repair in vivo

Finally, to determine whether enhancing lymphatic activation promotes skeletal regeneration *in vivo*, we evaluated local VEGFC delivery in a mouse tooth extraction model, a well-established model of alveolar bone injury and repair (Fig. 3d). Local administration of VEGFC-loaded hydrogel significantly increased LYVE1-positive lymphatic vessel density within the healing extraction socket, confirming enhanced lymphangiogenesis at the injury site (Fig. 3d). This was accompanied by significantly enhanced bone regeneration, demonstrated by increased bone volume fraction (BV/TV), trabecular number (Tb.N), and trabecular thickness (Tb.Th) measured by micro-computed tomography, together with increased bone formation detected by Masson’s trichrome staining (Fig. 3e). Consistent with enhanced osteogenesis, VEGFC treatment also increased expression of the osteogenic transcription factor Osterix (OSX), which regulates osteoblast differentiation, together with increased expression of BMP2, a key inducer of bone formation, and osteopontin (OPN), a marker of mature bone matrix deposition and mineralization (Fig. 3e). Further, we systemically administered VEGFC to mice for 3 weeks and found that, compared with PBS-treated controls, VEGFC-treated mice exhibited increased bone mineral density and Osterix expression across multiple skeletal sites, including femur, mandible, and vertebrae (Supplementary Fig. 1 and 2). Collectively, these findings demonstrate that enhancing bone lymphatic activation stimulates lymphangiogenesis, augments regenerative lymphangiocrine signaling, and accelerates bone regeneration following skeletal injury.

### Mandibular diseases are characterized by impaired lymphatic endothelial programs across transcriptomic, proteomic and genetic analyses

Given that bone lymphatic endothelial cells acquire a pro-regenerative phenotype during skeletal repair, we next asked whether disruption of lymphatic signaling represents a feature of mandibular diseases. We first analyzed human mandibular bone specimens obtained from healthy individuals and patients with chronic periodontitis using label-free quantitative proteomics (Fig. 4a). Consistent with the transcriptomic findings, periodontitis samples exhibited significant reductions in proteins associated with lymphatic endothelial identity and vascular guidance. In particular, the lymphatic marker LYVE1, together with the vascular guidance molecules EPHB4, NRP1, PLXND1, and PLXNA1, were all significantly decreased in diseased bone compared with healthy controls (Fig. 4a). The concordance between independent transcriptomic and proteomic datasets obtained from distinct mandibular diseases suggests that suppression of lymphatic-associated signaling represents a conserved feature of pathological bone remodeling within the craniofacial skeleton.

**Figure 4.**
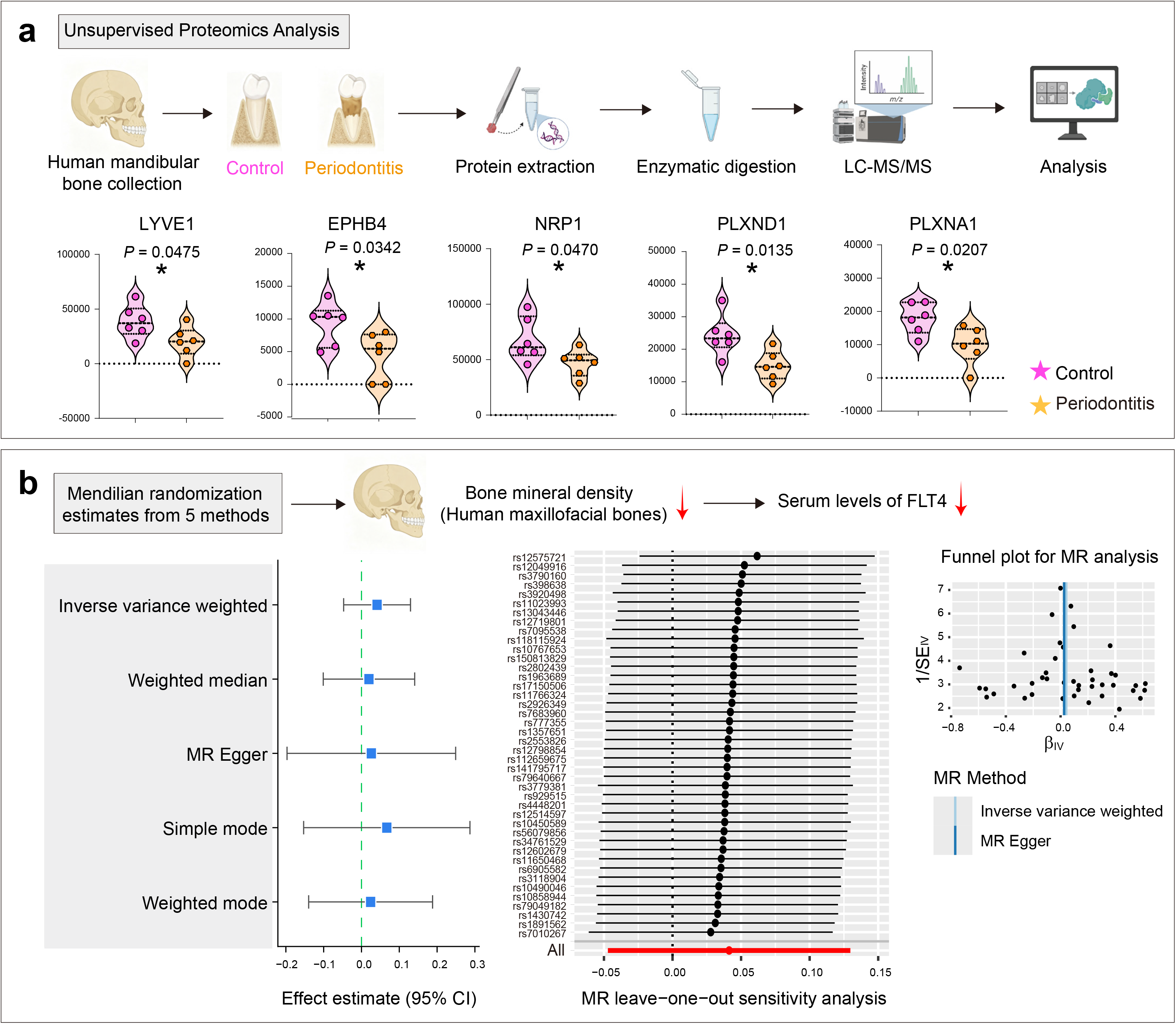
Lymphatic impairment in human maxillofacial bones during mandibular diseases. **a** Proteomics analysis of human mandibular bone samples from control and periodontitis groups. Violin plots showing differential expression of LYVE1, EPHB4, NRP1, PLXND1, and PLXNA1 between control and periodontitis groups. *P* values are derived from two-tailed unpaired t tests. * *P* < 0.05; *n=6*. **b** Two-sample Mendelian randomization analysis assessing the causal association between genetically predicted DXA-derived head bone mineral density and serum levels of the protein FLT4. Forest plots show MR estimates obtained using inverse-variance weighted (IVW), MR-Egger, weighted median, simple mode, and weighted mode methods. Leave-one-out analyses and funnel plots demonstrate sensitivity analyses assessing the robustness of the MR findings.

Next, we investigated whether lymphatic signaling is genetically associated with mandibular bone homeostasis by performing two-sample Mendelian randomization using genetically predicted head bone mineral density as the exposure and circulating FLT4 (VEGFR3) levels as the outcome (Fig. 4b). Across five independent Mendelian randomization approaches, including inverse-variance weighted, weighted median, weighted mode, simple mode, and MR-Egger analyses, genetically predicted reductions in head bone mineral density were consistently associated with lower serum FLT4 levels (Fig. 4b). Leave-one-out analyses demonstrated that the observed association was not driven by individual genetic variants, while funnel plot analyses showed no evidence of substantial directional pleiotropy (Fig. 4b), supporting the robustness of the findings. Collectively, these complementary transcriptomic, proteomic and genetic analyses consistently identify impaired lymphatic signaling as a hallmark of mandibular disease and implicate the VEGFC–FLT4 signaling axis as a fundamental regulator of maxillofacial bone homeostasis.

### Therapeutic activation of bone lymphatics restores bone repair in osteonecrosis of the jaw

The consistent reduction of lymphatic-associated genes and proteins across mandibular diseases suggested that impaired lymphangiogenesis may contribute directly to defective skeletal repair. We therefore asked whether therapeutic activation of bone lymphatics could restore tissue regeneration in bisphosphonate-related osteonecrosis of the jaw (BRONJ), a clinically challenging condition characterized by impaired vascularization and delayed healing following tooth extraction^57-60^.

To enhance local lymphangiogenesis, VEGFC-loaded hydrogels were implanted into extraction sockets immediately following tooth extraction in zoledronate-treated mice, and tissues were analyzed two weeks later (Fig. 5a). Three-dimensional immunofluorescence imaging demonstrated a striking increase in LYVE1-positive lymphatic vessels throughout the healing socket following VEGFC treatment compared with PBS-treated controls. Quantitative analysis confirmed a several-fold increase in lymphatic vascular density (Fig. 5a), demonstrating effective activation of local lymphangiogenesis. Importantly, VEGFC treatment simultaneously increased the abundance of Endomucin (EMCN)-positive blood vessels, indicating that stimulation of lymphatic growth was accompanied by enhanced angiogenesis within the regenerating extraction socket (Fig. 5a). These findings suggest that activation of bone lymphatics promotes coordinated regeneration of both lymphatic and blood vascular networks during mandibular repair.

**Figure 5.**
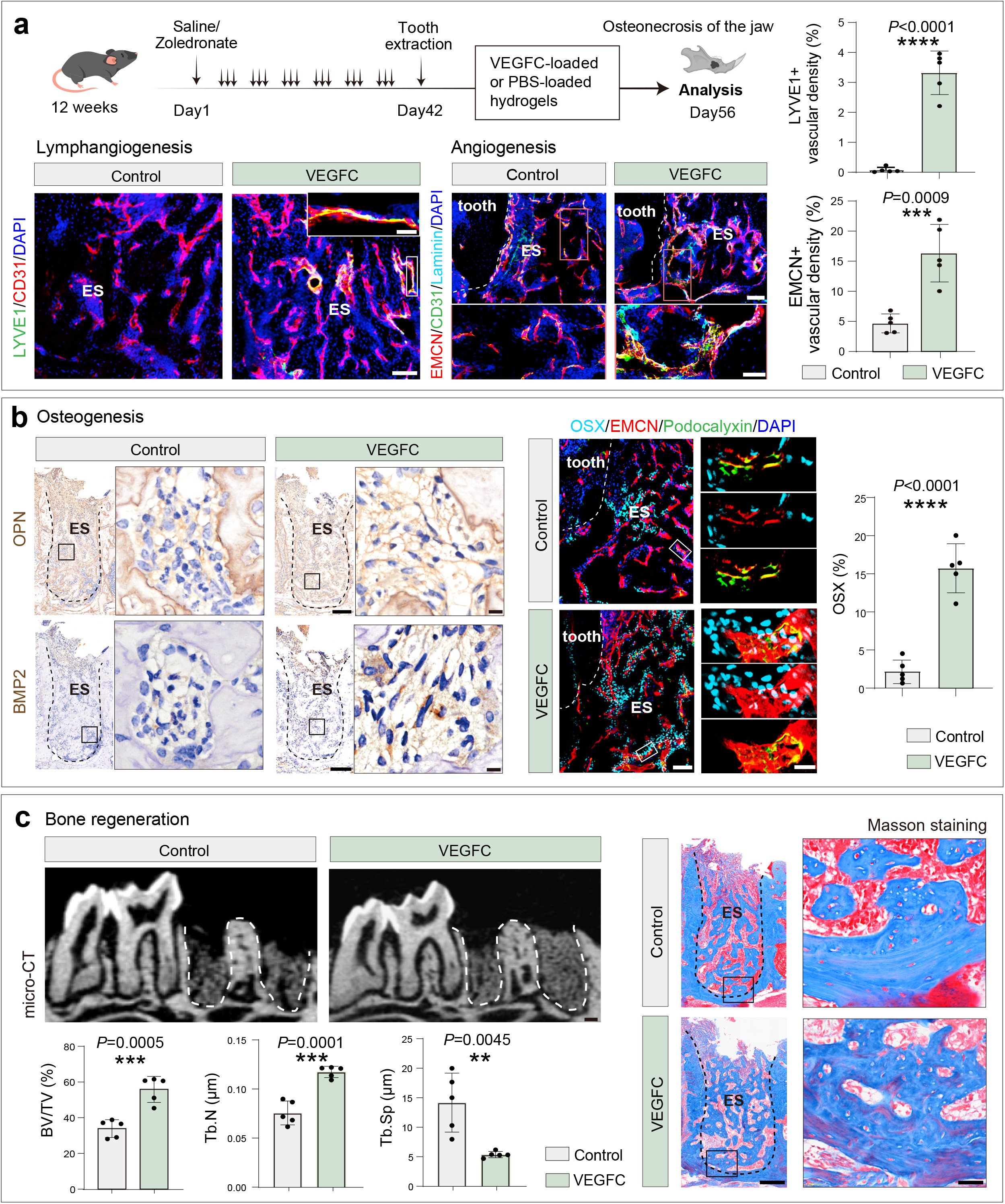
Bone lymphatics promote mandibular repair in bisphosphonate-related osteonecrosis of the jaw. **a** Schematic illustration of VEGFC-loaded hydrogel treatment in the bisphosphonate-related osteonecrosis of the jaw (BRONJ) model. Representative 3D images showing LYVE1, CD31, EMCN and Laminin immunostaining in control and VEGFC-treated groups. Bar graphs showing quantifications of LYVE1 and EMCN vascular density. *P* values are derived from two-tailed unpaired *t* tests. *** *P* < 0.001 and **** *P* < 0.0001; *n=5*. Scale bars: 100 μm (overview) and 20 μm (insets). **b** Representative images showing OPN and BMP2 immunohistochemical staining and OSX, EMCN, and Podocalyxin immunofluorescence staining. Bar graphs showing quantification of Osx (%). *P* values are derived from two-tailed unpaired *t* tests. **** *P* < 0.0001; *n=5*. Scale bars: for left panel, 150 μm (overview) and 40 μm (insets); for right panel, 100 μm (overview) and 10 μm (insets). **c** Representative micro-CT images and Masson staining images in control and VEGFC-treated groups. Bar graphs showing quantifications of micro-CT parameters. *P* values are derived from two-tailed unpaired *t* tests. ** *P* < 0.01 and *** *P* < 0.001; *n=5*. ES, extraction socket. Scale bars: for micro-CT, 200 μm; for Masson staining, 150 μm (overview) and 40 μm (insets).

Because successful vascular regeneration is closely coupled to osteogenesis, we next examined whether VEGFC-induced lymphatic activation enhanced bone formation. Immunohistochemical analyses demonstrated substantially increased expression of the osteogenic proteins osteopontin (OPN) and BMP2 within VEGFC-treated extraction sockets compared with controls (Fig. 5b). Consistent with these observations, immunofluorescence imaging revealed a marked increase in Osterix (OSX)-positive osteoprogenitor cells closely associated with regenerating EMCN-positive blood vessels, indicating enhanced osteoblast differentiation within the vascularized repair tissue (Fig. 5b). Further, osteoclasts in the extraction sockets remained unchanged following VEGFC treatment compared with controls (Supplementary Fig. 3), suggesting that VEGFC-induced lymphatic activation does not affect osteoclastogenesis. These findings demonstrate that restoration of lymphatic signaling promotes osteogenic differentiation in parallel with vascular regeneration.

We next assessed whether these cellular and molecular changes translated into improved structural regeneration of the jawbone. Micro-computed tomography demonstrated that VEGFC-treated mice exhibited significantly greater bone regeneration than controls, with increased bone volume fraction (BV/TV) and trabecular number (Tb.N), together with reduced trabecular separation (Tb.Sp), indicating restoration of trabecular architecture within the extraction socket (Fig. 5c). Histological evaluation using Masson’s trichrome staining further confirmed abundant deposition of newly formed collagen-rich bone matrix in VEGFC-treated animals compared with controls (Fig. 5c). Together, these findings demonstrate that therapeutic activation of bone lymphatics restores lymphangiogenesis, promotes angiogenesis and osteogenic differentiation, and ultimately enhances functional bone regeneration in experimental ONJ (Fig. 6).

**Figure 6.**
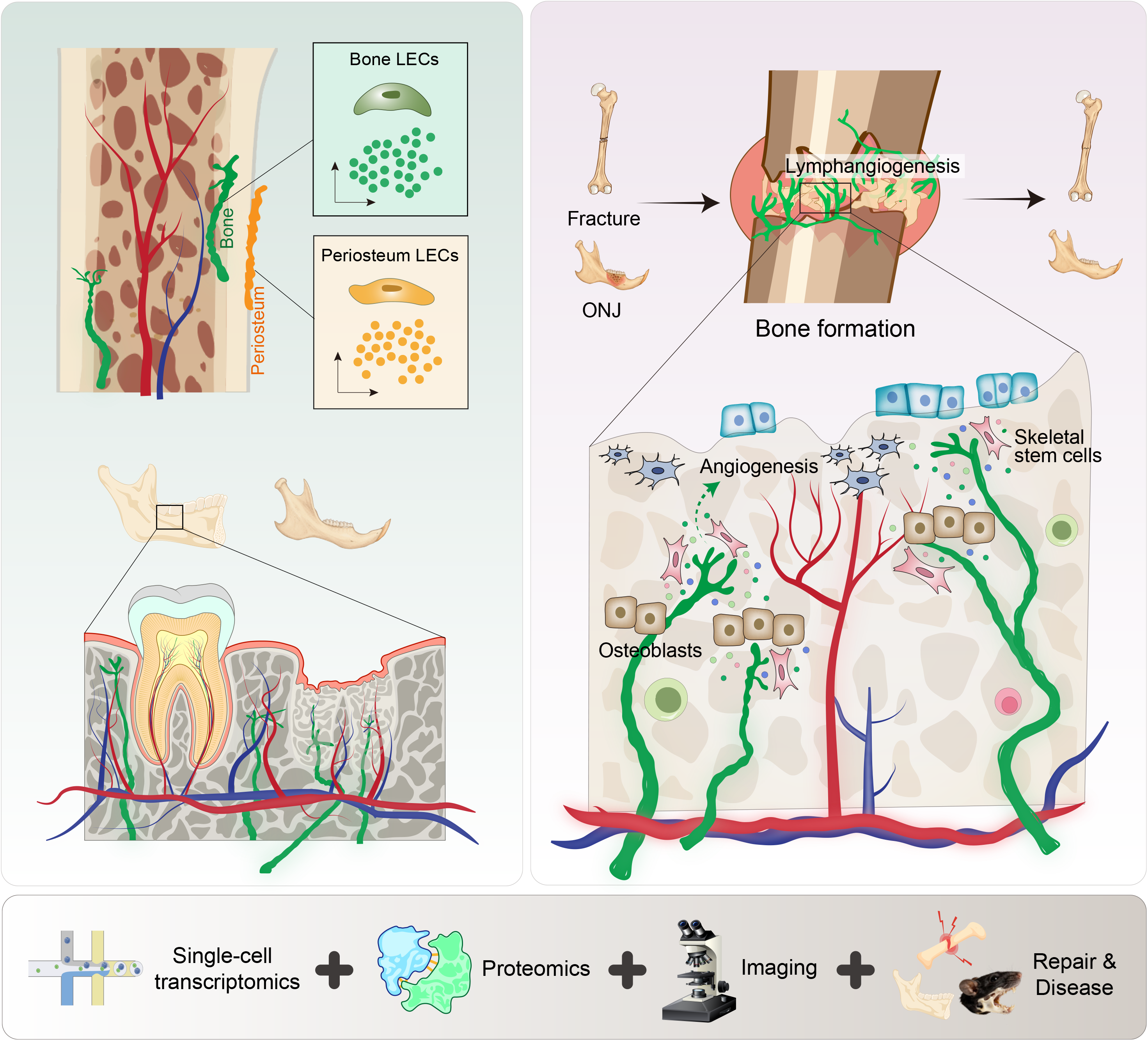
Transcriptomic, proteomic and spatial evidence for lymphatic-driven bone repair and regeneration

## Discussion

Our study identifies bone lymphatics as an essential component of the mandibular and long bone regenerative microenvironment and demonstrates that they actively drive bone repair. By integrating single-cell transcriptomics, proteomics, imaging and functional studies, we show that bone lymphatic endothelial cells acquire a regenerative program during repair and can be therapeutically targeted to restore skeletal regeneration.

A key finding of this study is that bone lymphatics are molecularly and functionally distinct from periosteal lymphatics. Our analyses consistently separated bone and periosteal endothelial populations, demonstrating that bone lymphatics possess a distinct transcriptional identity. Further our findings support the existence of a specialized bone lymphatic niche and demonstrate that periosteal lymphatics cannot fully account for the molecular signatures or regenerative functions of lymphatic endothelial cells within bone.

Our multi-omics analyses further demonstrate that impaired lymphatic signaling is a common feature of periodontitis and osteonecrosis of the jaw. Despite their distinct etiologies, these disorders converge on suppression of lymphatic-associated pathways, suggesting that disruption of the lymphatic niche represents a shared mechanism contributing to impaired mandibular regeneration. Importantly, therapeutic activation of the VEGFC–FLT4 axis restored lymphangiogenesis, enhanced osteogenesis, and significantly improved bone regeneration in experimental osteonecrosis of the jaw. These findings identify lymphatic signaling as a therapeutically actionable pathway and suggest that restoring lymphatic function may represent a regenerative strategy not only for ONJ but also for other mandibular disorders characterized by impaired bone healing, including traumatic bone defects and delayed fracture repair.

More broadly, our findings add to the emerging paradigm that lymphatic vessels are positive regulators of bone regeneration and repair. Together with recent studies demonstrating lymphangiocrine regulation of regeneration in multiple organs^22,24,25,61^, our work advances bone lymphatics as a fundamental component of the skeletal regenerative niche. Beyond redefining skeletal vascular biology, these findings provide a conceptual framework for developing lymphatic-based regenerative therapies that complement existing osteogenic approaches. Targeting and activation of the lymphatic vasculature may therefore offer new therapeutic opportunities to enhance bone repair, improve craniofacial reconstruction, accelerate healing following dental and maxillofacial surgery, and treat challenging skeletal diseases in which conventional regenerative strategies remain inadequate.

## Methods

### Mice

C57BL/6J mice (8-12 weeks old) were purchased from Dashuo Biotechnology Co., Ltd. (Chengdu, China). All animal experiments complied with local ethical guidelines and regulations and received approval from the Research Ethics Committee of West China Hospital of Stomatology (Approval No. WCHSIRB-D-2023-245).

### Murine tooth extraction model and VEGFC hydrogel injection

12-week-old wild-type C57BL/6 mice were anesthetized, and their bilateral mandibular first molars were minimally invasively extracted. Mice with root fractures were strictly excluded from the study. Immediately post-extraction, the sockets were injected with dextran methacrylate (DexMA; EFL, EFL-DexMA-500K) hydrogels loaded with recombinant human VEGFC protein (20 ng/mL; Novoprotein, C546) or PBS in control mice. At 2 weeks post-surgery, the mice were euthanized, and the harvested mandibles were fixed in 4% paraformaldehyde overnight at 4°C.

### Bisphosphonate-related osteonecrosis of the jaw (BRONJ) model

Wild-type C57BL/6 mice were randomly assigned to saline control or zoledronate treatment groups. Mice in the BRONJ group received intraperitoneal injections of zoledronate (200 μg/kg; MCE, HY-13777) from day 1 to day 42, whereas control mice received an equivalent volume of sterile saline. On day 42, bilateral mandibular first molars were extracted, and the extraction sockets were treated with DexMA hydrogels containing VEGFC (20 ng/mL) or PBS. After achieving complete hemostasis, the wounds were sutured. All mice were euthanized on day 56 (14 days after extraction) for subsequent tissue collection.

### Systemic VEGFC administration

Eight-week-old mice were intraperitoneally injected daily with VEGFC protein (100 ng/g body weight/day; Novoprotein, C546) or an equivalent volume of PBS for 21 consecutive days. Mandibles, femurs, and spines collected from all experimental groups were fixed in 4% paraformaldehyde overnight at 4°C.

### micro-CT

Mandibular samples were scanned using μCT45 micro-CT scanner (SCANCO) with a voxel size of 10.0 μm, 55 kVp, and 145 μA. Three-dimensional rendering was performed using CTvox software (Bruker). Quantitative 3D morphometric parameters, including bone volume fraction (BV/TV), trabecular number (Tb.N), and trabecular thickness (Tb.Th), were analyzed utilizing CTAn software (Bruker).

### Masson’s trichrome staining

Samples were decalcified in 0.5 M EDTA (pH 7.4) for 2 days, dehydrated through a graded ethanol series, and embedded in paraffin. Sagittal sections (5 μm thickness) were prepared, deparaffinized, and rehydrated for subsequent staining. Masson’s trichrome staining was performed using a commercial staining kit (Solarbio, G1340) according to the manufacturer’s instructions. Stained sections were imaged using a digital slide scanner (SLIDEVIEW VS200, Olympus).

### Immunohistochemistry

Paraffin sections were deparaffinized, rehydrated, and subjected to antigen retrieval in EDTA buffer (pH 9.0). After blocking, sections were incubated overnight at 4°C with primary antibodies against BMP2 (Affinity Biosciences, AF5163), OPN (Proteintech, 22952-1-AP), and IL-1β (Servicebio, GB11113), followed by incubation with HRP-conjugated goat anti-rabbit IgG (H+L) secondary antibody (Servicebio, GB23303). Immunoreactivity was visualized using DAB, followed by hematoxylin counterstaining. Images were acquired using a slide scanner (VS200, Olympus).

### Immunofluorescence imaging

Freshly dissected bone samples were carefully cleared of muscles, periosteum, and surrounding soft tissues, followed by fixation in 4% paraformaldehyde (PFA) on ice for 2.5 h. Samples were decalcified in 0.5 M EDTA at room temperature for 2 days, with the decalcification solution replaced twice daily, and subsequently washed three times with PBS. Decalcified specimens were cryoprotected in 20% sucrose (Sigma-Aldrich, V900116) containing 2% polyvinylpyrrolidone (PVP; Sigma-Aldrich, P5288) at 4 °C for 12 h. Samples were then embedded in 8% gelatin supplemented with 20% sucrose and 2% PVP and sectioned at 70–100 μm thickness using a Leica CM1950 cryostat (Leica Microsystems, Germany). Following rehydration, sections were permeabilized with 0.3% Triton X-100 and blocked with 5% donkey serum. Primary antibodies (see Supplementary Table 1 for detailed information) were incubated with sections at 1:150 dilution for 4 h at room temperature, followed by incubation with Alexa Fluor-conjugated secondary antibodies (1:300 dilution) for 1–1.5 h. Nuclei were counterstained with DAPI (Solarbio, C0060), and sections were mounted with Fluoromount-G (Invitrogen, 00-4958-02) and stored at 4 °C until imaging. Fluorescence imaging was performed using a spinning-disk confocal microscope (IXplore IX83 SpinSR, Olympus). Image processing and three-dimensional reconstruction were carried out using Imaris software (version 10.2.0).

### TRAP staining

Tartrate-resistant acid phosphatase (TRAP) staining was performed on paraffin-embedded sagittal sections according to the manufacturer’s instructions (Solarbio, G1492). Briefly, sections were deparaffinized and rehydrated, followed by incubation with the TRAP staining solution. After staining, sections were counterstained, dehydrated, and mounted for imaging. TRAP-positive multinucleated cells were identified as osteoclasts. All stained sections were digitized using a whole-slide scanning system (VS200, Olympus).

### Cell culture

Human lymphatic endothelial cells (hLECs, IM-H480, IMMOCELL) were cultured in Endothelial Cell Medium (ECM, CM-H026, Procell) at 37°C with 5% CO_2_. Human bone marrow mesenchymal stem cells (hBMSCs, CP-H166, Procell) were cultured in α-MEM (Gibco, 12492013) supplemented with fetal bovine serum (FBS; Sigma-Aldrich, F7524) under standard culture conditions at 37°C with 5% CO_2_.

### Scratch assay

hLECs and HUVECs were seeded into 6-well plates and cultured to confluence, followed by serum starvation for 12 h. A scratch wound was generated using a sterile pipette tip, and cells were treated with VEGFC (20 ng/mL) or an equal volume of PBS in low-serum medium. Wound closure was imaged by phase-contrast microscopy at 0, 6, and 12 h.

### Transwell non-contact co-culture system

A paracrine co-culture system was established using 0.4 μm pore Transwell inserts (Corning). hBMSCs were seeded in the lower chambers and cultured in osteogenic induction medium containing dexamethasone (100 nM), β-glycerophosphate (10 mM), and ascorbic acid (50 μg/mL). LECs were seeded in the upper chambers and treated with recombinant VEGFC (20 ng/mL) or PBS.

### Alkaline phosphatase and Alizarin Red S staining

Bone MSCs were assessed for osteogenic differentiation by alkaline phosphatase (ALP) and Alizarin Red S (ARS) staining on days 7 and 14. For ALP staining, cells were fixed with 4% PFA and stained using a BCIP/NBT Color Development Kit (Beyotime, C3206). ALP activity was quantified using an ALP Assay Kit (Beyotime, P0321S), according to the manufacturer’s instructions. For ARS staining, cells were fixed and stained with OriCell^®^ ARS solution (Cyagen Biosciences, ALIR-10001) to visualize mineralized nodules. Quantitative analysis was performed by eluting bound ARS with 10% cetylpyridinium chloride (CPC) for 30 min at 37°C and measuring absorbance at 562 nm.

### Mendelian randomization analysis

The potential effect of genetic liability to DXA-derived head bone mineral density on serum levels of the protein FLT4 was considered using a two-sample Mendelian randomization (MR) approach. Genetic variants associated with head bone mineral density at genome-wide significance (*P* < 5×10^-8^) were selected as instrumental variables. To ensure the independence of genetic instruments, we performed linkage disequilibrium (LD) clumping using information from the 1000 Genomes Project European ancestry population, applying a ±10 Mb genomic window and an LD r² < 0.001. Causal estimates were obtained using five complementary MR methods, including inverse-variance weighted (IVW), MR-Egger, weighted median, simple mode, and weighted mode approaches. Sensitivity analyses, including leave-one-out analysis and funnel plot assessment, were performed to evaluate the robustness of the findings.

### Single cell RNA-sequencing data analysis

scRNA-seq datasets were retrieved from publicly available repositories, namely-Gene Expression Omnibus (GEO) and BioStudies (Supplementary Table 2 and 3). For high-level scRNA-seq processing, Seurat R toolkit 5.4.01^62^ was employed in R version 4.5.2. Seurat objects for samples of interest were created with cells having >500 unique genes each expressed in a minimum of 3 cells. For downstream analysis, cells having <800 UMIs, more than the sample-specific 98th percentile of UMIs, >6,000 detected genes, or >10% mitochondrial transcripts were filtered out. Lastly, scDblFinder^63^ package was used to identify and remove potential doublets.

After merging filtered cells from all samples, data normalization was performed on the merged object using *NormalizeData* function (normalisation method=“LogNormalize”; scaling factor=10,000). Top 2,000 variable features were computed using *FindVariableFeatures* function (method=“variance stabilizing transformation”). Genes related to cell cycle (GO:0007049) were removed from the identified variable features. Using these filtered variable genes, data was scaled with *ScaleData* function. Principal component analysis (PCA) was performed with *RunPCA* function, followed by selection of reasonable principal components guided by elbow plots (*ElbowPlot*). Further, batch effects were corrected with the help of Harmony (v1.2.3)^64^ using *RunHarmony* function using sample identity as the covariate and selected components. Further, non-linear dimensionality reduction was performed with *RunUMAP* function on the Harmony-corrected embeddings. Finally, graph-based clustering following “Louvian algorithm” was carried out with *FindNeighbors* and *FindClusters* functions on optimum resolution. Cell clusters were manually annotated based on expression of canonical markers (Supplementary Table 4), visualized with *FeaturePlot*, *VlnPlot*, and *DotPlot* functions.

Differentially expressed genes (DEGs) were identified using *FindMarkers* function (min.pct=0.1, test.use=“wilcox”). Genes with an adjusted p-value < 0.05 and average log_2_(fold change) > 0.5 were considered significantly upregulated.

Gene Ontology (GO) term enrichment (ont=“BP”, pAdjustMethod=“BH”) was carried on significantly upregulated genes using clusterProfiler^65^ package and org.Mm.eg.db database with p-and q-value cutoff = 0.05.

For GO term scoring, respective genes were obtained from MSigDB R database (category=“C5”, subcategory=“BP”). Scores were computed as the average normalized expression of genes and visualized with DotPlot.

Group-specific average normalized expression values of selected genes were used to generate a gene-by-group expression matrix visualized in the form of heatmap generated using the *pheatmap* R package.

PCA was performed on the scaled average normalized expression matrix of highly variable genes using *prcomp* function. Sample scores for the first two principal components were used to generate PCA plots.

For hierarchical clustering analysis, pairwise sample similarity of average normalized expression matrices of highly variable genes was assessed using Spearman’s rank correlation, which was converted to a distance matrix (1−correlation) to generate a dendrogram using *hclust* function (method=“complete”).

CellChat^66^ package was employed to infer cell-cell communication across conditions using the normalized gene expression matrices. *netVisual_circle* function was used to visualize interactions of cell type of interest. Interactions outgoing from cell type of interest were filtered and grouped based on targets to compute target-specific number of ligand–receptor interactions and the total communication strength. Interaction summaries were visualized as bubble plots, where dot size represented the number of interactions and dot colour represented the total interaction strength.

For secretome analysis, genes expressing secreted proteins were identified among DEGs using secreted protein database obtained from UniProt (organism_id: 10090; cc_scl_term: SL-0243) and further GO term enrichment was performed.

Code used for all the analysis is made available in Supplementary Information.

### Human alveolar bone specimen collection

All human sample collection procedures were approved by the Research Ethics Committee of West China Hospital of Stomatology, and written informed consent was obtained from all participants. Human alveolar bone specimens were obtained from male and female donors in the periodontitis and healthy control groups. The donors ranged in age from 42 to 56 years. The chronic periodontitis group consisted of patients with moderate-to-severe periodontitis characterized by clinical attachment loss, alveolar bone resorption, and tooth extraction due to irreversible periodontal destruction. The healthy control group included individuals with clinically healthy periodontal tissues undergoing tooth extraction for orthodontic treatment or impacted teeth. Participants with systemic metabolic disorders, autoimmune diseases, malignancies, long-term corticosteroid or immunosuppressive therapy, acute oral infections, or a history of head and neck radiotherapy were excluded. Following extraction, alveolar bone specimens were immediately rinsed with pre-chilled PBS on ice to remove residual blood and granulation tissue, transferred into pre-chilled cryotubes, and stored in liquid nitrogen until proteomic analysis.

### Proteomic analysis

Frozen human alveolar bone samples were pulverized in liquid nitrogen, and total proteins were extracted using a bone-specific lysis buffer supplemented with protease inhibitors. Protein extraction was followed by ultrasonication and centrifugation, and protein concentrations were determined using a BCA assay. Protein quality was assessed by SDS-PAGE. Qualified samples underwent reduction, alkylation, and tryptic digestion, followed by desalting and lyophilization. The resulting peptides were analyzed by label-free quantitative LC-MS/MS. Raw data were processed against the human reference proteome database for protein identification and quantification. Differentially expressed proteins between periodontitis and healthy controls were identified and subjected to functional enrichment analyses.

### Statistical analysis

Statistical analyses were performed using GraphPad Prism (version 9.0). No statistical methods were used to predetermine sample sizes. Mice were randomly allocated to experimental groups, and samples were processed in an arbitrary order. No blinding was performed, and no animals or samples were excluded from the analyses. Data are presented as mean ± SD. Comparisons between two groups were performed using two-tailed unpaired Student’s *t*-tests unless otherwise indicated. For single-cell RNA sequencing data, differential gene expression analyses were performed using the Wilcoxon rank-sum test. Statistical significance was defined as *P* < 0.05. Significance is denoted as: * for *P* < 0.05, ** for *P* < 0.01, *** for *P* < 0.001, **** for *P* < 0.0001

## Data availability

The proteomic raw data generated in this study have been deposited in the iProX repository under accession number IPX0018626000.

## Acknowledgments

A.P.K. is supported by Ministry of Education (MOE) Singapore: Academic Research Funds (#024983-00001, #025277-00026), European Research Council (StG: metaNiche, 805201), and European Union’s Horizon 2020 (857524). J.C. is supported by the National Natural Science Foundation of China (82422021, 82270961), Sichuan Provincial Health Commission (24QNMP015), and West China Hospital of Stomatology Sichuan University (RD-03-202401, RCDWJS2024-5). This work is supported by the National Institutes of Health (NIH). M.V.R. is supported by the National Institute on Aging (R01AG073349) and National Institute of Arthritis and Musculoskeletal and Skin Diseases (R01AR055655 and R01AR082460). S. J. is supported by the Nanyang Technological University Research Scholarship (Reg. No. 200604393R). Y.Y. is supported by National Nature Science Foundations of China (No. 824B2026).

## Contributions

S.J., J.L., J.Z., and H.C. methodology, investigation, and visualization. Y.W. and M.Y. investigation. Y.Y. analysis. M.V.R. reviewing and editing. J.C. writing, validation, and formal analysis. A.P.K. writing, methodology, supervision, reviewing, and editing.

---

**Supplementary Figure 1. VEGFC administration increases bone mass across multiple skeletal sites. a** Schematic overview of systemic VEGFC administration via intraperitoneal route in 8-week-old mice. **b** Representative micro-CT reconstruction images of the mandible in control and VEGFC-treated groups. Bar graphs showing quantifications of micro-CT parameters. *P* values are derived from two-tailed unpaired *t* tests. ** *P* < 0.001; *n=5*. Scale bars: 200 μm (overview) and 100 μm (insets). **c** Representative micro-CT reconstruction images of the femur. Scale bars: 200 μm (overview) and 75 μm (insets). *P* values are derived from two-tailed unpaired *t* tests. * *P* < 0.05; ** *P* < 0.001; *n=5*. Scale bars: 200 μm (overview) and 75 μm (insets). Representative 3D images showing immunolabeling for OSX and EMCN in the femur. Nuclei: DAPI. Scale bars: 100 μm (overview) and 20 μm (insets).

**Supplementary Figure 2. VEGFC administration increases bone mass in vertebra. a** Schematic overview of systemic VEGFC administration via intraperitoneal route in 8-week-old mice. **b** Representative micro-CT reconstruction images of the L4 lumbar vertebra in control and VEGFC-treated groups. *P* values are derived from two-tailed unpaired *t* tests. ** *P* < 0.001 and *** *P* < 0.0001; *n=5*. Scale bars: 100 μm. Representative 3D images showing immunolabeling for OSX and EMCN in vertebra. Nuclei: DAPI. Scale bars: 100 μm (overview) and 50 μm (insets).

**Supplementary Figure 3. Bone lymphatics do not alter osteoclastogenesis in BRONJ. a** Representative images showing TRAP staining of the first molar extraction socket in the bisphosphonate-related osteonecrosis of the jaw (BRONJ) model. Scale bars: 400 μm (overview) and 20 μm (insets).

**Supplementary Table 1** Antibody list

**Supplementary Table 2** List of datasets and specific samples used for cross-tissue LEC analyses

**Supplementary Table 3** List of datasets and specific samples used for LEC analyses in control vs fracture conditions

**Supplementary Table 4** List of canonical markers used to annotate cell clusters

